# Aerobic anoxygenic phototrophic bacteria are ubiquitous in phyllo- and endosphere microbiomes of boreal and subarctic plants

**DOI:** 10.1101/2023.02.19.529139

**Authors:** Riitta Nissinen, Ole Franz, Salla Kovanen, Meri Mäkelä, Venla Kraft, Katri Ketola, Alli Liukkonen, Kati Heikkilä-Huhta, Heikki Häkkänen, Janne A. Ihalainen

## Abstract

In addition to oxygenic photosynthetic systems, solar radiation is utilized for energy by diverse anoxygenic photosynthetic systems. Aerobic anoxygenic phototrophic bacteria (AAPB) perform photosynthesis without producing oxygen but still live in aerobic conditions. Typically they have been reported from aquatic ecosystems, but they can also be found from polar and desert soil ecosystems and primary succession communities like soil crusts. Recently, AAPB have been discovered in the metagenomic data of several plant phyllospheres. By utilizing citizen science, we screened plant foliar samples from eleven different locations in Finland for AAPB by near infrared fluorescence imaging of culturable phyllosphere and endosphere bacteria. Near infrared fluorescence reports the presence of AAPB which contain Bacteriochlorophyll *a* molecules, embedded in Light Harvesting 1 - Reaction Center complex. We found that AAPB were ubiquotous in phyllosphere communities of diverse plant species in all sampling locations. They were also consistently present in the endosphere of plant species with perennial leaves. Most of the AAPB isolated represent alphaproteobacterial genera Sphingomonas and Methylobacterium, but several isolates from genus Lichenihabitans as well as putative novel alphaproteobacterial taxa were also identified. Methylobacterial isolates were mostly detected in the phyllosphere with weak host specificity, while Sphingomonas AAPB were detected also in the endosphere of several plant species, with clear host specific taxa. We studied also the fluorescence spectral properties of several AAPBs. All the observed spectra resemble typical fluorescence spectral properties of Light Harvesting complex 1. Still, slight variation among each spectra could be obtained, revealing some physical difference among the complexes. Our results demonstrate for the first time, that AAPB are common in cold climate plant endophytic as well as epiphytic microbiomes and they build up substantial amounts of Bacteriochlorophyll *a* containing Light Harvesting complexes. Their putative role in plant adaptation to strong seasonality in light and temperature or tolerance of abiotic stressors remains to be investigated in future studies.

## I. INTRODUCTION

All major ecosystems on our planet depend on plants, algae and cyanobacteria, as they are able to convert energy from light into organic compounds. In addition to these organisms, photosynthetic systems are also widely present in evolutionary ancient anoxygenic phototrophic bacteria, including aerobic anoxygenic phototrophic bacteria (AAPB)^1^. By using Bacteriochlorophyll a (BChl *a*) molecules, AAPB are able to utilize solar radiation to produce energy in the form of ATP. Typically, they contain type II reaction center and LH1 Light-harvesting complexes^2^, which typically form LH1-RC supercomplexes^3^. AAPB do not fix carbon, and thus depend on organic carbon for growth. In contrast with purple sulfur bacteria, AAPB are aerobic, and the photosynthesis is not inhibited by oxygen. However, these bacteria do not produce oxygen, as instead of using water, these bacteria rely on diverse organic compounds as electron donors for photosynthesis^4^.

AAPB have been originally discovered in oligotrophic marine environments^1^, and are ubiquotous in both marine and freshwater ecosystems^5,6^. In aquatic ecosystems, AAPB can constitute up to a third of total bacterial populations. They are considered to play a significant role in carbon transformations in marine ecosystems^4^. Photosynthesis has been speculated to supply these phototrophic organisms supplemental energy to enable utilization of recalcitrant organic compounds in low carbon environments. However, environmental factors impacting AAPB abundance, diversity and their photosynthesis are still relatively poorly understood.

AAPB were long thought to be mainly restricted to aquatic ecosystems. However, in the last decade, AAP-associated genes and AAPB have been discovered in several terrestrial ecosystems, including polar and glacier soils^7–9^, and soil crusts^10,11^. AAPB have been detected also in plant phyllosphere (surfaces of plant aboveground tissues) microbiomes by metagenome mining^12,13^, and by culturomics^14^. Discoveries of AAPB in association with plants are particularly intriguing, as microbes have been shown to be crucial for several aspects of plants’ wellbeing, including growth and development, as well as resistance to biotic and abiotic stressors, including drought, pollutants, heat and low temperatures^15,16^. Plant associated microbes need to adapt to life in planta and light is known to impact interactions of plants with pathogens as well as with commensal and beneficial bacteria^17–19^. Yet, comprehensive surveys of AAPB presence in plant phyllosphere or endosphere communities are not available, and their diversity, biogeography and putative interaction with plants remain to be characterized.

Here, we present results of systematic screening and characterization of culturable AAPB in leaf phyllo- and endosphere of multiple plant species in ten distinct sampling locations across latitudal gradient spanning 1000 km and five climate zones in Finland. We show, that AAPB are consistently present in plant phyllosphere microbiomes, as they are detected virtually in all plant species and in locations from oroarctic to hemiboreal climate regions. Additionally, we report for the first time, that AAPB are also present as endophytes of several perennial plants. Further, all our spectrally analyzed plant associated AAPs show fluorescence properties of LH1-RC complexes with small variation of spectral shapes, which indicates variation in the pigment organization in the LH1 complexes.

## II. RESULTS

### A. Detection of AAPB in phyllo- and endosphere samples

We screened the phyllospheres and endospheres of 23 boreal and arctic plant species from total of ten geographically distinct locations in Finland (Figure 1 and Table I) for presence of AAPB by near infrared (NIR)-fluorescence imaging of culturable bacteria. The NIR region of radiation relates to BChl *a* fluorescence emission which is indicative for AAP (Supplemental Figure S1). 12 plant species were screened in replicated manner (3-24 replicate samples, Table I). For 11 additional plant species, one replicate per species was analyzed. After incubation for seven weeks at +4°C, AAP positive bacterial colonies were detected in cultures from all 12 plant species tested with replicated sampling, and in five out of 11 species with single sampling (Supplemental Table 1).

**FIG. 1:**
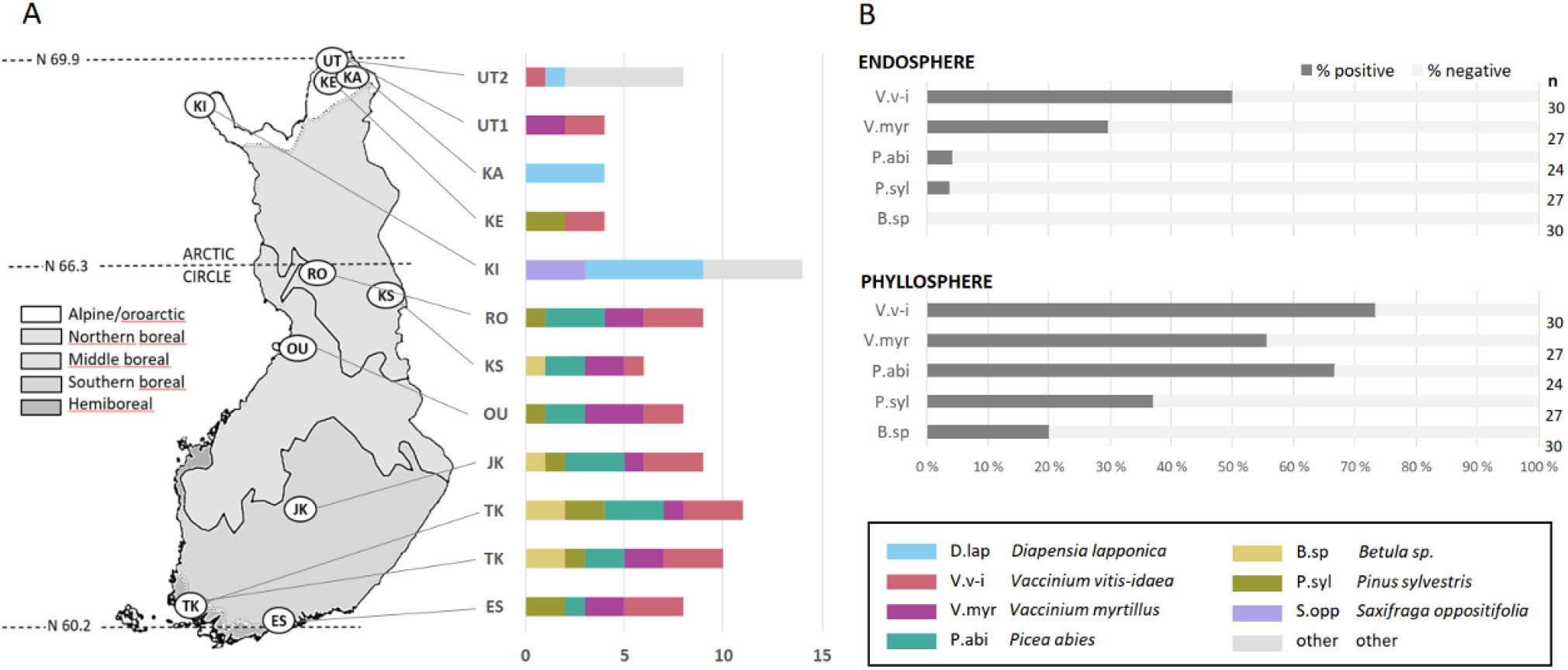
Sampling sites, collected plant species, and proportion of AAP positive samples. A) Locations and climate regions of the sampling sites with AAP positive plant samples. The bars represent the number of AAP positive plant samples in each site, with the plant species indicated by different colors. The detailed information of species and sample numbers are tabulated in (Table I). Plant species other than the core set of the species of our study *(Betula* spp, *Pinus sylvestris, Picea abies, Vaccinium vitis-idaea,* and *Vaccinium myrtillus*) were collected only from the UT2, KA and KI sampling sites. B) The proportions of the AAP positive and negative endosphere and phyllosphere samples of the five core plant species in this study.

**Table I:** The sampling locations, with coordinates, climate zone, the plant species and number of biological replicates collected from each site.

| Sampling location | coordinates | climate zone | species sampled | nr of replicates |
| --- | --- | --- | --- | --- |
| Utsjoki 2 (UT2) | 69.91, 27.06 | oroarctic | C, D.lap, E, I, J, K, Q, V.v-i, WB, WG, WH, WL, WQ, WS | 1 |
| Utsjoki 1 (UT1) | 69.89, 27.01 | northern boreal | B.sp, V.myr, V.v-i | 3 |
| Kaldoaivi (KA) | 69.82, 27.50 | oroarctic | D.lap | 4 |
| Kevo (KE) | 69.75, 27.01 | northern boreal | B.sp, P.syl, V.v-i | 3 |
| Kilpisjärvi (KI) | 69.01, 20.50 | oroarctic | B.sp, D.lap*, H, N, L, Q, S.opp | 3 (6*) |
| Rovaniemi (RO) | 66.30, 25.72 | northern boreal | B.sp, P.abi, P.syl, V.myr, V.v-i | 3 |
| Kuusamo (KS) | 65.57, 29.84 | northern boreal | B.sp, P.abi, P.syl, V.myr, V.v-i | 3 |
| Oulu (OU) | 65.02, 25.46 | middle boreal | B.sp, P.abi, P.syl, V.myr, V.v-i | 3 |
| Jyväskylä (JK) | 62.14, 25.45 | southern boreal | B.sp, P.abi, P.syl, V.myr, V.v-i | 3 |
| Turku (TK) | 60.28, 22.19 | hemiboreal | B.sp, P.abi, P.syl, V.myr, V.v-i | 6 |
| Espoo (ES) | 60.11, 24.45 | southern boreal | B.sp, P.abi, P.syl, V.myr, V.v-i | 3 |

Plant species: B.sp: *Betula pendula*/*B. pubescens*, C: *Kalmia procumbens*, D.lap: *Diapensia lapponica*, E: *Phyllodoce caerulea*, H: *Oxyria digyna*, I: *Vaccinium uliginosum*, J: *Deschampsia flexuosa*, K: *Spinulum annotinum* ssp. *alpestre*, L: *Dryas octopetala*, N: *Bistorta vivipara*, P.syl: *Pinus sylvestris*, P.abi: *Picea abies*, Q: *Betula nana*, WB: *Empetrum nigrum* ssp., WG: *Salix* sp., hermafroditum, WS: *Marchantiophyta* sp., S.opp: *Saxifraga oppositifolia*, V.v-i: *Vaccinium vitis-idaea*, V.myr: *Vaccinium myrtillus*, WH: *Arctous alpina*, WL: *Huperzia selago*, WQ: *Polytrichum* sp

AAPB were present in the phyllosphere samples of all AAP positive plant species (17/23). Additionally, AAPB were also detected in the leaf endosphere samples of seven plant species: *Diapensia lapponica, Saxifraga oppositifolia, Dryas octopetala, Vaccinium vitis-idaea, Vaccinium myrtillus, Pinus sylvestris* and *Picea abies*. In plant species with more than three samples analyzed, AAPB were present in 20-100 %and 0-91 %of the samples from phyllosphere and endosphere, respectively (Figure1).

AAPB were most common in the endospheres of *Diapensia lapponica* and *Vaccinium vitis-idaea* with NIR-fluorescence positive colonies detected in 11/11 and 22/30 endophere samples analyzed, respectively. Further, AAPB were isolated from 3/3 and 2/3 endosphere samples from *S. oppositifolia* and *D. octopetala*, respectively (Figure 1, Table I). The lowest occurrence rates of AAPB were in Betula samples, where no AAP positive colonies were detected in any of the 30 leaf endosphere samples analyzed, and only 6 out of 30 phyllosphere samples were positive.

AAPB were present in samples from all sampling locations used in the study, ranging from 60.2°N to 69.9°N, and from hemitemperate to oroarctic climate (Figure 1, Table I), with no marked differences in proportion of positive samples between locations.

### B. Taxonomic distribution, host and niche affiliation of plant associated AAP bacteria

Of the total of over 2000 AAP positive colonies detected, we purified and identified 88 representative isolates by partial 16S rRNA sequencing. Most of the sequenced isolates belong to bacterial class Alphaproteobacterial families Sphingomonadales and Rhizobiales (Figure 2). Majority of those represent genera Sphingomonas (49 isolates) and Methylobacterium (34 isolates). Additionally, four isolates closely related to Lichenibacterium (Lichenihabitans) within family Rhizobiales, and one isolate closely related to genus Aurantimonas (Aurantimonadaceae, order Hyphomicrobiales) were detected (Figure 2).

**FIG. 2:**
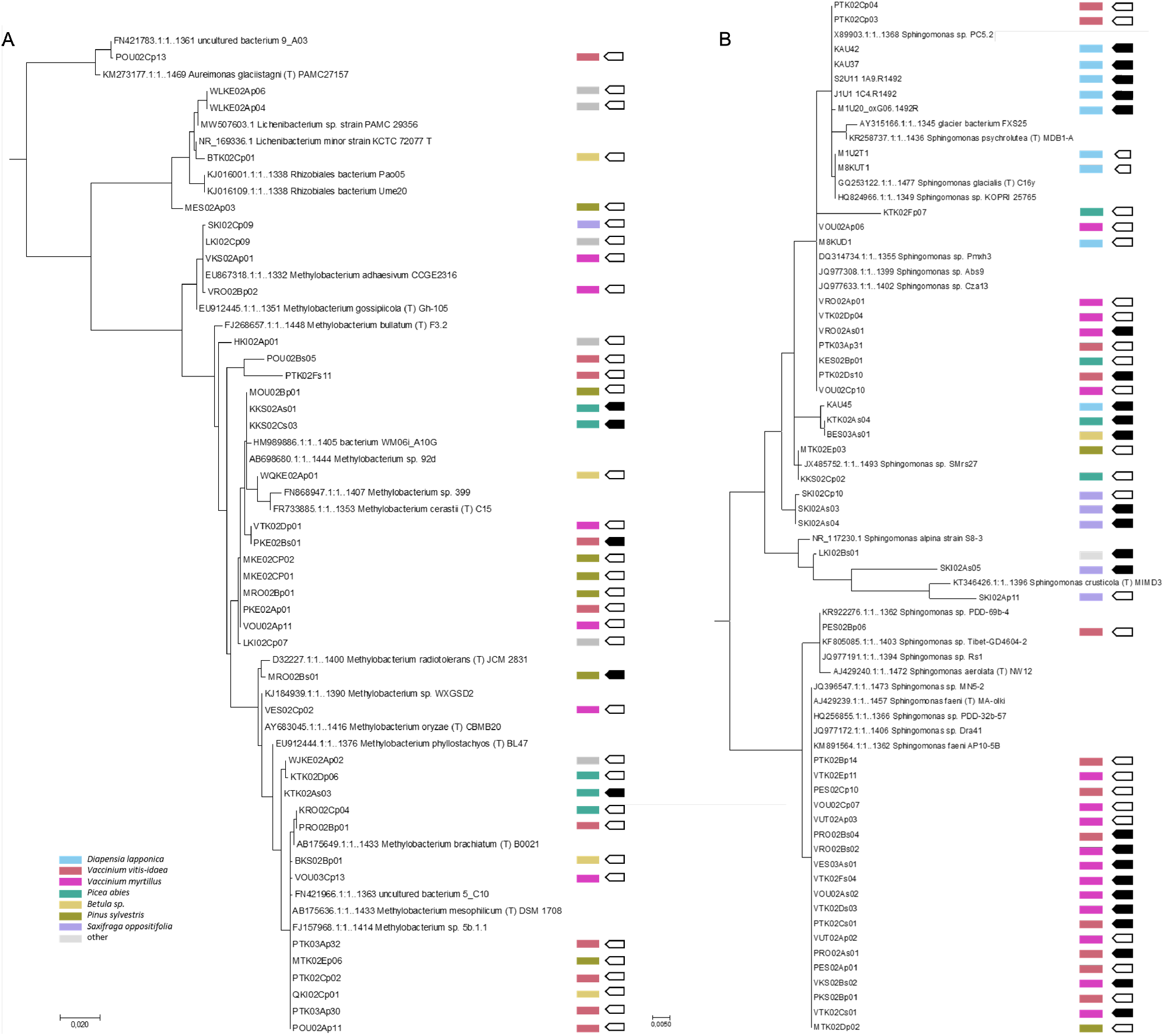
Phylogenetic trees of AAP positive isolates. A) Rhizobiales isolates, with sequences from most similar type strains or isolates included B) Sphingomonas isolates with sequences from most similar type strains or isolates from public databases. The colored blocks correspond to the host plant of the strains, open arrows correspond to epiphytic, and filled arrows correspond to endophytic bacteria. The coding of the samples reflect the geographic location and time period of the year sample collected (See Table I and Figure 1) Maximum likelihood trees were constructed using MEGA v 11

A clear majority of the 49 positive Sphingomonas strains clustered with species *Sphingomonas faeni* or *Sphingomonas glacialis*. Strains clustering with *S. faeni* were almost exclusively detected in *Vaccinium myrtillus* and *V. vitis-idaea*, with only one isolate originating from *P. abies*. These strains were detected in both phyllosphere and endosphere samples of the two Vaccinium species, in which they were detected in all sampling locations (Figure 1).

Most of the strains with high 16S rRNA gene sequence identity to *S. glacialis* were isolated from *Diapensia lapponica* (8 strains), *Vaccinium vitis-idaea* (4 strains), *Vaccinium myrtillus* (5 strains) and *Picea abies* (2 strains). These strains were also repeatedly isolated from endosphere as well as phyllosphere of their host plants (Figure 2).

Unlike Sphingomonas strains, AAP positive Methylobacterium isolates represent diverse Methylobacterium species, and were detected in many different plant species, with no clear host plant affiliation. AAP positive Methylobacterium strains were isolated most often from *P. sylvestris, P. abies, V. vitis-idaea* and *V. myrtillus*, and clear majority originated from phyllosphere samples. Only five Methylobacterium strains out of the total 34 identified were isolated from endosphere samples. Out of these five, three isolates originated from *P. abies* endosphere, one from *V.-vitis-idaea*, and one from *P. sylvestris*.

In addition to AAP strains clustering with Methylobacterium species, four isolates with less clear taxonomic status in order Rhizobiales were isolated. Of the publicly available bacterial 16S rRNA sequences, most similar sequences originate from bacteria from diverse lichens in Antarctica and in boreal Russian Karelia, including strains proposed as type strains for novel genus Lichenibacterium^20,21^.

One strain, isolated from *V. vitis-idaea* phyllosphere, was classified as *Aureimonas* sp., with closest matching sequence from an uncultured bacterial clone from soybean phyllosphere (accession FN421783) (Figure 2).

### C. Fluorescence spectral analysis of the selected AAP positive bacteria

AAP positive bacteria were identified by fluorescence imaging through a band pass filter which allows detection in the spectral region around 880 nm, as described in Supp Fig **S1**. We confirmed BChl *a* production, and the type of the photosystem by measuring fluorescence spectra from representative bacterial isolates (Figure 3). An excitation wavelength of around 390 nm, which excite the Soret band of BChl *a* molecules, turned out to be much more effective excitation wavelength than the 400 – 500 nm region. This is in line with that at least the Sphingomonadales contain a large amount of carotenoids which are not involved in excitation energy transfer to the BChl *a* molecules^22,23^

**FIG. 3:**
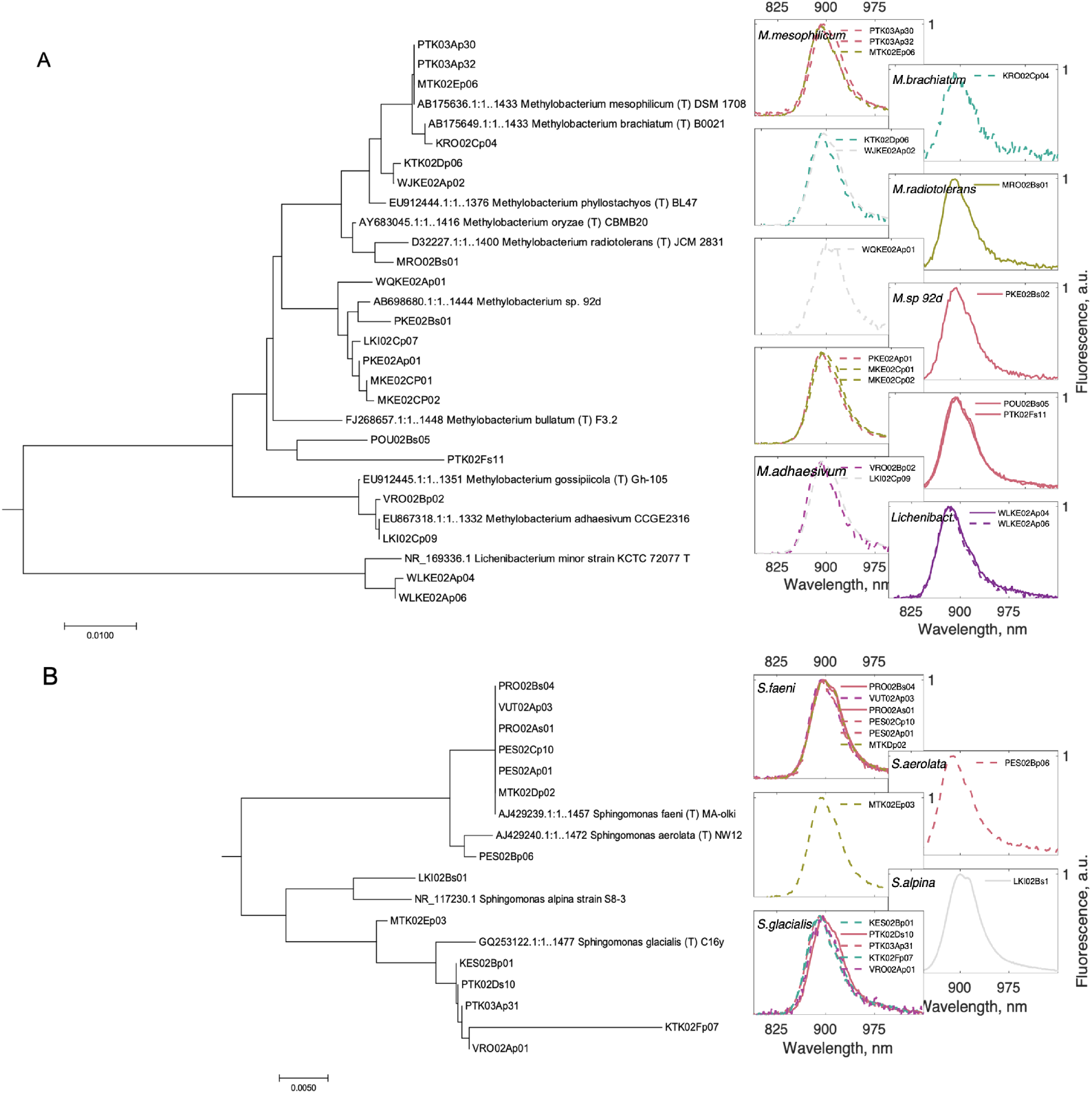
Phylogenetic trees of AAP positive isolates with selected fluorescence spectra. A compressed phylogenetic tree (from Figure 2) with their corresponding fluorescence emission spectra. A)Rhizobiales bacteria and B)Sphingomondales type of bacteria. The color coding indicates the plant host as in Figure 2. The solid lines correspond to endophytic bacteria and dashed lines to phyllospheric bacteria. The sample coding is described in Table I)

By spectral analysis we could confirm, that the fluorescence detected by imaging indeed originates from BChl *a* containing complexes in all inspected isolates. The spectra show emission features in the region of 900 nm, which imply that the strains in this study carry complexes containing LH1-RC-complex only, typical for AAPBs^1,24^. The spectral position was also similar to what was detected in Sphingomonas sp. AAP5^25^. A closer look to the spectral shapes reveals, however, an interesting phenomenon of the spectral properties of the samples. The precise fluorescence maxima, and generally, the spectral shapes varied between different samples. The spectral properties were not directly dependent on bacterial or plant species or on sampling location.

The emission features of all samples can be divided in three different spectral regions. The blue most emission band is located between 875 nm and 885 nm, the middle one at around 895 nm and the red most band between 900 nm and 905 nm. The relative proportion of each these bands varies between the species but also between isolates of the same species collected from different locations or from different plant species. In some of the samples, such as *Methylobacterium brachiatum* or *Sphingomonas faeni* the blue most emission is located around 875 nm.

A slight trend with Sphingomonas samples can be seen that the endophytic samples show generally more redshifted maximum of the fluorescence spectra in comparison to the phyllospheric samples. This, however, remains to be confirmed with a larger data set in the future studies.

## III. DISCUSSION

AAPB have long been associated mainly with marine and freshwater ecosystems, where they can constitute up to 30 %of total bacterial communities^1,4^. Recently, AAPB have been reported also from terrestrial habitats, including plant phyllospheres, but the abundance and prevalence in plants has thus far remained uncharted. In this study, we show, that AAPB are ubiquitous in plant phyllosphere, and frequently present also in endophytic plant microbiomes in boreal and arctic ecosystems. Our findings strongly suggest, that plant phyllosphere as well as endosphere - globally largest biologically active surfaces - constitute a major, thus far understudied habitat for anoxygenic photosynthesis, at least in the strongly seasonal cold ecosystems targeted in this study. Previously AAPB or AAP associated genes have been described from phyllosphere communities of soybean, clover, Arabidopsis and wheat in Central Europe^12,14^ and from agaves and cacti in deserts^13^, indicating, that these bacteria seem to be present in phylloshere communities in diverse ecosystems.

We detected AAPB in phyllosphere communities of all plant species sampled in a replicated manner, and in all sampling locations spanning different climate zones. Taking into account, that the core plant species sampled - lingonberry, bilberry, spruce, pine and birch are key species in boreal ecosystems, we can conclude, that AAPB represent a major functional group in boreal and subarctic phytobiomes.

A vast majority of the AAP positive isolates from phyllosphere in this study are bacteria from alphaproteobacterial taxa Methylobacterium and Sphingomonas. Our findings largely corroborate previous reports, where these two genera, in particular Methylobacterium, were the main taxa in phyllosphere AAPB communities of temperate climate and desert plants^12–14,26^. Methylobacterium and Sphingomonas are both well known as core members of phyllosphere plant microbiomes of diverse plants, and they are metabolically well suited for phyllospheric lifestyles, as both are able to utilize plant metabolites as their carbon source^17^. The ability to utilize solar radiation as an energy source likely provides these bacteria an additional adaptation mechanism to oligotrophic and radiation rich phyllosphere environment. Sphingomonas strains isolated from phyllosphere have been shown to protect their host plant from invasion by plant pathogenic bacteria by nutritional niche occupancy and via plant immune priming^27,28^. This plant protective ability was specific to plant associated strains (Sphingomonas isolates from other environmental sources were not effective), highlighting the importance of adaptation to plant associated lifestyle, although AAP capacity of the strains was not investigated in these studies^27,28^. Additionally, several strains from novel bacterial genus Lichenihabitans (Lichenibacterium) were detected in the phyllosphere samples of birch, pine and northern clubmoss (*Huperzia selago*). Intriguingly, this bacterial genus was recently describe as a core member of large set of lichen microbiomes globally^29^.

In addition to phyllosphere, we detected consistent presence of AAPB in the leaf endospheres of diverse plant species, mainly in perennial leaves or photosynthetic tissues. Thus far, anoxygenic photosynthesis has been only reported from plant phyllosphere, with the exception of nitrogen fixing photosynthetic Bradyrhizobial strains forming specialized stem nodules in select legumenous plant species^30^.

Most of the endophytic AAPB in this study represent genus Sphingomonas. Unlike highly diverse AAP positive Methylobacterial isolates with no clear host plant association, most Sphingomonas isolates in this study clustered closely with two species, *S. glacialis* and *S. faeni*, and these endophytic strains had restricted host plant range. *S. glacialis* strains were detected predominantly in *Diapensia lapponica*, a hemiarctic plant species, that is extremely cold tolerant and highly resistant to drought and solar radiation. Intriguingly, *S. glacialis* has been shown to be part of Diapensia’s core microbiome, being consistently present in the plant’s endo- and epiphytic communities, including plant seeds^31^ (Nissinen unpublished). Further, all *S. glacialis* strains from *D. lapponica* have been shown to be AAP positive (Nissinen, unpublished), as are *S. glacialis* isolates described from an arctic glacier cryoconite^32^ and from an alpine high elevation lake^25^.

Strains from this study clustering with *S. faeni* were present almost exclusively in leaves and photosynthetic stem tissues of two perennial shrub species *V. vitis-idaea* and *V. myrtillus,* and were detected in these plants very frequently, in all sampling locations and in high relative abundances, suggesting a very tight association between photosynthetic *S. faeni* and these host plants. *S. faeni* type strain has been described from straw material from Finland^33^, but the type strain genome does not contain any genes indicative for AAP. Other *S. faeni* strains with high sequence similarity to isolates in this study originate from high alpine plants from China^34^, as well as from diverse plant phyllospheres, antarctic lichen communities, leaf tissues of an arctic plant *Oxyria digyna*^31,35^, but also from microbial communities of clouds^36^. The AAP status of these strains have not been examined.

AAPs contain BChl *a* molecules which absorb light at different wavelengths from chloroplasts in plants. As previously reported for AAPs, the BChl *a* molecules are embedded in RC-LH1 complexes^1,5,30,37^. However, in few rare cases LH2 complexes have been found from such bacteria^38^. So far, in our fluorescence spectroscopic analysis we found only RC-LH1 complexes. It is well known that the spectral shapes are extremely sensitive to the properties of the LH1 complexes, where small structural changes of the protein complex lead to changes in the excitonic manifold of the system, and therefore changes in fluorescence spectral shapes^39^. In our fluorescence experiments, we failed to find clear correlations between the sample species and fluorescence character. The blue most emission bands are rather blue-shifted from the main emission bands. One option for such blue shift of emission band could be that some of the LH1 complexes are constructed from Zn-chelated BChl *a* molecules, which show about seven nanometer blue shifts in absorption^24^. This, however, remains to be confirmed in the pigment analysis of the bacteria.

It is notable, that although the spectra were measured at rather similar time points after harvesting, the growth state of strains may be different due to different growth rate of the bacteria. It is, however, unlikely that each bacterial species would have single type of fluorescence emission spectral properties and the species annotation would become possible via the fluorescence spectral analysis. Still, general spectral trends should be collected for larger data sets to be more conclusive in this respect.

Most of plant microbiome research has thus far focused on root and rhizosphere microbial communities. Far less is known about microbiota in the photosynthetic plant tissues. Plant aboveground tissues are challenging environments as microbial habitats. Yet, our understanding on bacterial adaptations to leaf environments are still limited. Phyllosphere is highly oligotrophic, and phyllospheric microbes are exposed to radiation and strongly fluctuating temperature and humidity^40–43^. In the phyllosphere, AAP could be beneficial by providing energy for utilization of recalcitrant plant metabolites and for phyllosphere competence.

In contrast, plant endospheres are not limited for plant derived carbon and energy. The consistent presence of AAPB in leaf endosphere of several species addressed in this study, and their vertical transmission^31^ suggests an alternative role for anoxygenic photosynthesis for endophytes. In case of photoactive Bradyrhizobium strain ORS278, forming stem nodules in Aeschynomene species, AAP genes are expressed in stem nodules, and are required for efficient stem nodulation, nitrogen fixation and growth enhancement of their host plant, providing evidence role of anoxygenic photosynthesis in plant-microbe symbiosis^30^. Intriguingly, the AAPB identified in this study represent well known plant associated bacterial taxa, some with proven status as member of host plant core microbiome^31^. However, further studies are needed for dissecting the putative role of AAP in plant-microbe interactions for broader group of leaf endophytic AAPB.

## IV. METHODS

### A. Sampling

The majority of the samples were collected as citizen science project in collaboration with seven high schools (Utsjoen saamelaislukio, Rovaniemen Lyseon lukio, Kuusamon lukio, Oulun Steiner lukio, Jyväskylän lyseon lukio, Otaniemen lukio and Turun Kerttulin lukio) in Finland during late summer 2022. Additionally, in this study, isolates and samples collected from Kevo, Kilpisjärvi and Kaldoaivi regions in Finnish Lapland (years 2014-2022) were used. The sampling sites, representing five different climate regions from hemiboreal to oroarctic climate, are indicated in Figure 1a. Detailed information on sampling sites, plant species and number of samples collected and sampling time(s) are in Supplemental Table 1. The students collected the plant samples aseptically, following standardized protocol and using sterile tools, gloves and collection boxes, and shipped the samples to laboratory on ice, where they were processed within 72 hours.

### B. Sample processing and bacterial isolation

Plant material was trimmed to remove any dead tissue, and leaves or needles were picked aseptically using sterile tools. Phyllosphere microbes were removed by ultrasonication in sterile 20 mM potassium phosphate buffer, pH 6.5 (KPi) with 0.015% Silwet-L77 for 3 min. Epiphytes were pelleted by centrifugation (13 000 g, 3 min) of 2×1 ml of the supernatant, and the pellets were resuspended in total 200 μl of KPi. Plant material was surface sterilized by immersion into 3% NaOCl (3 min), followed by triple rinse in sterile distilled water (3 × 1 min). Sterility checks were performed by plating 100 μl of last rinsing water on R2A plates. Sterilized material was homogenized in sterile stomacher bags in KPi (5 mL buffer per 1g of plant tissue). The macerate was used for serial dilution plating on 50% strength R2A solid medium with pH adjusted to 6.5 (Difco). The plates were incubated at 23°C for 3 days, transferred to 4°C and screened for BChl *a* containing colonies 2, 4 and 6 weeks after plating (See description below and Fig **S1**). BChl *a* positive colonies were transferred to new plates and re-analysed for Bchl *a* after 10 days (3 days RT, 1 week at 4°C). Individual colonies were picked to produce pure cultures, which were preserved in R2B, pH 6.5 with 30% glycerol at −80°C.

Selected isolates were assigned to bacterial taxa by amplification with primers 27F and 1492R^44^ and subsequent sequencing (primer 1492R) of bacterial 16S rRNA genes. For sequence alignment, Ribosomal Database Project 16S rRNA training set 18 was used (http://rdp.cme.msu.edu/). For phylogenetic analyses, isolate sequences were aligned with sequences from the closest bacterial type strains and other relevant bacterial sequences using MEGA (Molecular Evolutionary Genetics Analysis) version 11^45^. 16S rRNA sequences have been submitted to NCBI with accession numbers xx-xx.

### C. Imaging of the Petri dishes

The Petri dish imaging system combines a LED excitation either at visible region or at 395 nm for white light imaging or for NIR-fluorescence imaging, respectively. The NIR-fluorescence emission light was detected by using a band pass filter with maximum wavelength at 880 nm and with FWHM 70 nm (Thorlabs). The images (white light for visualization of all bacterial colonies on Petri dish or NIR-limited image for visualization of BChl *a* containing bacteria) were taken with Rasperry Pi camera. (Franz, unpublished). The setup was a compact version based on^46^.

### D. Fluorescence emission spectroscopic analysis

The bacterial colonies were excited with laser light at 390 nm laser light directly from the Petri dishes. The position of the excitation light was adjusted with help of white light imaging and a web camera and blocked afterwards. The fluorescence light was guided to a spectrograph (Acton 300) attached with NIR-array detector (Andor). The spectral calibration was performed by adjusting the pixel values with the emission peaks from the Hg(Ar) calibration lamp. The spectra were not corrected for the detector sensitivity, but the dark noise and background were subtracted from the measured spectra.

## V. ACKNOWLEDGMENTS

The participating high schools classes and their teachers from Utsjoen saamelaislukio, Rovaniemen Lyseon lukio, Kuusamon lukio, Oulun Steiner lukio, Jyväskylän lyseon lukio, Otaniemen lukio and Turun Kerttulin lukio, are highly appreciated for collecting the majority of the plant samples for this study. The financial support for RN and JAI from KONE foundation is gratefully acknowledged. The authors also wish to thank the personnel at Kilpisjärvi research station (University of Helsinki) and Kevo subarctic research station (University of Turku) for their help in sampling, and Martin Chilman for his helpful suggestions for improving the figures of the paper.

---

**FIG. S1:**
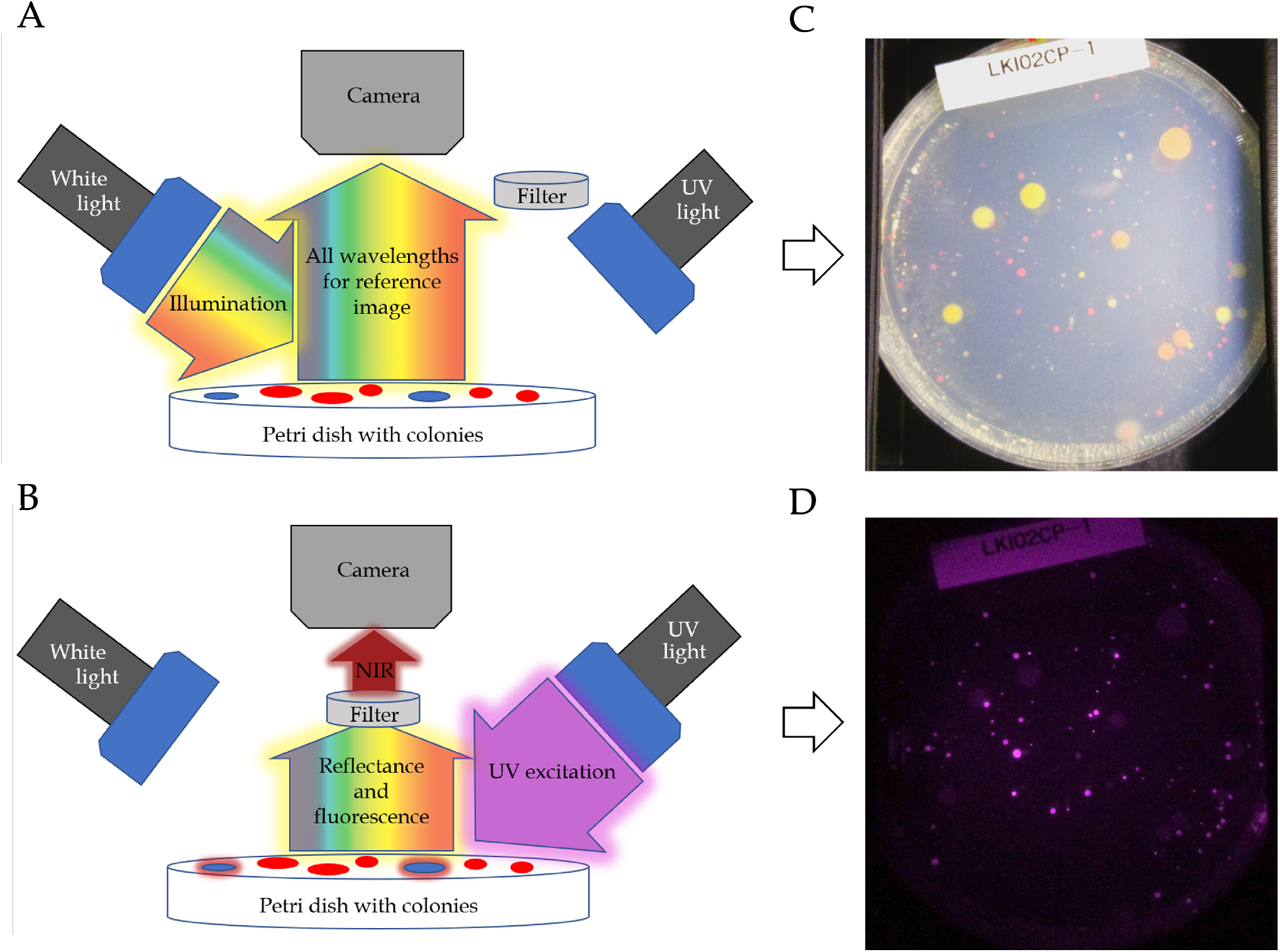
Working principle of the NIR-imaging device. A) White light imaging. Petri dishes are illuminated with white light and the picture of the dish is captured with a Rasberry Pi PiNoir Camer V2 with ordinary RGB-settings. B) UV-excitation induces fluorescence of the BChl *a* molecules in the samples and fluorescence is detected through a band pass filter, centered at 880 nm (FWHM 70nm, Thorlabs) by Rasberry Pi camera. C) An example of a white light image. D) An example of the fluorescence image from the same Petri dish. As the sample were kept still between the imaging processes, the images were superimposed, positive colonies were identified and collected for further analysis.

